# Mathematical Models for the Influence of Cytarabine on White Blood Cell Dynamics in Acute Myeloid Leukemia

**DOI:** 10.1101/428326

**Authors:** Felix Jost, Enrico Schalk, Kristine Rinke, Thomas Fischer, Sebastian Sager

## Abstract

We investigate the personalisation and prediction accuracy of mathematical models for white blood cell (WBC) count dynamics during consolidation treatment using intermediate or high-dose cytarabine (Ara-C) in acute myeloid leukemia (AML). Ara-C is the clinically most relevant cytotoxic agent for AML treatment.

We extend the gold-standard model of myelosuppression and a pharmacokinetic model of Ara-C with different hypotheses of Ara-C’s pharmacodynamic effects. We cross-validate 12 mathematical models using dense WBC count measurements from 23 AML patients. Surprisingly, the prediction accuracies are similarly good despite different modelling hypotheses. Therefore, we compare average clinical and calculated WBC recovery times for different Ara-C schedules as a successful methodology for model discrimination. As a result, a new hypothesis of a secondary pharmacodynamic effect on the proliferation rate seems plausible. Furthermore, we demonstrate how personalized predictions of the impact of treatment timing on subsequent nadir values could be used for clinical decision support.

**Author summary:** The major obstacle in accurately predicting the outcome of a medical therapy is the vast variation in individual response patterns. It concerns both the subjective experience of the patient and the objectively measurable achievement of a clinical remission with restoration of normal blood counts. Here, we address acute myeloid leukemia (AML)-chemotherapy using cytarabine (Ara-C) as this drug is this most important component of AML-treatment. In addition to the wide spectrum of genetic aberrations involved in pathogenesis leading to variations in patient response patterns, another facet of personalised medicine awaits exploration of its full potential: a systematic, mathematical approach to understand and manipulate the dynamics of relevant biomarkers. We use personalised mathematical models to describe and predict white blood cell (WBC) counts during AML consolidation treatment. We analyse why and to what extent low WBC counts, a serious adverse event during therapy, occur. In a comprehensive approach we investigate published models, compare them with our extended models and outline the impact of modelling assumptions and varying chemotherapy schedules on prediction accuracy and model discrimination. Our numerical results confirm the clinical finding that a newly proposed schedule is superior with respect to WBC recovery and shed new light on the reasons why.

## Preamble on terminology and potentially confusing synonyms

Our work is located in the intersection of mathematics, control theory, systems biology, pharmacology, and medicine. Words like “model” or “parameter” have different meanings in these scientific communities, and similar concepts have different names like “calibration”, “estimation”, or “personalisation”. For convenience, we list some synonyms that we did (not) use in **Table S1**.

## Introduction

Acute myeloid leukemia (AML) is a malignant clonal disorder of myeloid stem and progenitor cells. In untreated AML, immature neoplastic myeloid blasts rapidly proliferate and suppress the generation and maturation of blood cells in the bone marrow. While being a curable disease using chemotherapy including anthracyclines and/or cytarabine (Ara-C), this approach leads to prolonged myelosuppression with extremely low white blood cell (WBC) counts (leukopenia), i.e. values below 1 *G/L*, associated with a high risk of infection and treatment-related mortality [1].

Consolidation treatment, repetitive (up to 4) cycles of intermediate-/high-dose Ara-C (1 – 3*g/m^2^*) [2], is given once patients achieve complete remission (CR) and is considered the most important part of chemotherapy in preventing relapses. The current standard treatment of 3*g*/*m*^2^ Ara-C infusion lasting 3 hours every 12 hours on days 1, 3 and 5 for patients aged 60 years and younger was established by Mayer *et al*. [3].

If predictions from personalised mathematical models were reliable and accurate, they could be used for providing better care to AML patients receiving Ara-C consolidation treatment, e.g. in an automatized measurement–decision support loop [4, 5]. Precisely identifying the period of Ara-C-induced profound leukopenia and modification of treatment schedules based on such predictions might enable prevention of severe infectious complications, sepsis, and thus delay to undergo subsequent treatment cycles. Therefore, by realising timely adherence to consolidation therapy cycles and by avoiding delays in treatment schedule, the density of chemotherapy cycles may be increased and thus deeper remissions and lower relapse rates may be achieved. This may ultimately translate into improved overall survival rates.

Mathematical models for myelosuppression due to various chemotherapy agents have been proposed [6–11] and applied successfully to predict the dynamics of neutrophils [4, 12]. However, this is not the fact for high-dose Ara-C, the most important component in consolidation therapy [2, 13]. Pharmacology of Ara-C is particularly difficult, as its exact mechanisms of action both on normal and leukemic cells are not fully understood. The main effect of Ara-C on normal and leukemic proliferating cells is the inclusion of intracellular Ara-C triphosphate (Ara-CTP) into DNA and RNA, which impairs cell replication [14]. Yet, the synthesis of intracellular Ara-CTP is saturable such that the clinical success of intermediate-/high-dose Ara-C is not well explained [3, 15, 16]. Additional effects are the subject of ongoing research [16, 17].

Here, we surveyed different published and new hypotheses of the pharmacodynamic (PD) effects of Ara-C on WBC dynamics during AML consolidation therapy. We used models for myelosuppression and Ara-C pharmacokinetics (PK) from the literature to quantify prediction accuracies. The general modelling goals were to include possible secondary effects of Ara-C and to obtain a good balance between modelling detail, prediction accuracy, and the number of patient-specific parameters. As a successful methodology, we considered predictions of WBC recovery times (defined as the time when the WBC count recovers above 1*G/L*) for *different* Ara-C schedules and compared them to published average WBC recovery times.

## Methods

There are many different levels on which hematopoiesis and dynamics of leukemic cells can be modelled [10, 18–20]. We analysed models that capture only the most important dynamics for non-leukemic cells and “agglomerate” different physiological effects into simplifying expressions. Leukemic cell dynamics are not considered because of the lack of events (no measurable leukemic cells are present) and because of our focus on WBC recovery. The necessary model extensions to analyse relapse of leukemic cells are also beyond the scope of this paper. All model variations are based on the gold-standard model for myelosuppression developed by Friberg *et al*. [6] and are tailored to the special case of Ara-C via a parameterised two-compartment PK model. The personalised mathematical models (PMs) were generated by estimating model parameters from clinically measured WBC counts. To get a better understanding of the relation between modelling hypotheses and individual predictions, we used a point estimator approach. We also provide results of a population based estimation for verification and reference.

The procedure is described in detail below. The mathematical approaches to parameter estimation, uncertainty quantification, and statistical analysis, the mathematical equations, the different initial condition strategies, the PK model and the nonlinear mixed-effects modelling approach are described in **Appendix S1**.

### Mathematical Models

In the interest of providing a solid mathematical basis for future extensions we focused on the most important dynamical features. There are interesting recent extensions and ideas as possible additional effects of Ara-C [16, 17], inclusion of growth factors [1], or modelling also leukemic stem cells [20]. Higher levels of detail come at the price of an increased number of model parameters, which have to be estimated to obtain PMs. The current lack of clinical measurements of leukemic cells or drug and growth factor concentrations leads to identifiability issues with these additional model parameters. Thus we concentrated on agglomerating effects of Ara-C on proliferation and maturation rates.

**Fig 1** illustrates the basic assumptions from which we derived twelve model variations of the original Friberg model which we denote by M1–M12 from now on. They differ concerning the number of transition compartments (M1–M3), initial conditions for the differential equations (M3–M5), and model assumptions for the possible effects of Ara-C on proliferation and maturation rates (M5–M12). After intermediate evaluations of accuracies we concentrated on the most promising choice of scaling, transition compartments, and initial conditions, and included different modelling assumptions in the models M6–M12 which are alternatives to M5, our Ara-C extension of the gold-standard model for myelosuppression [6]. Most models refer to previous approaches in the literature and are included for a comprehensive comparison and evaluation of our new hypotheses.

**Fig 1.**
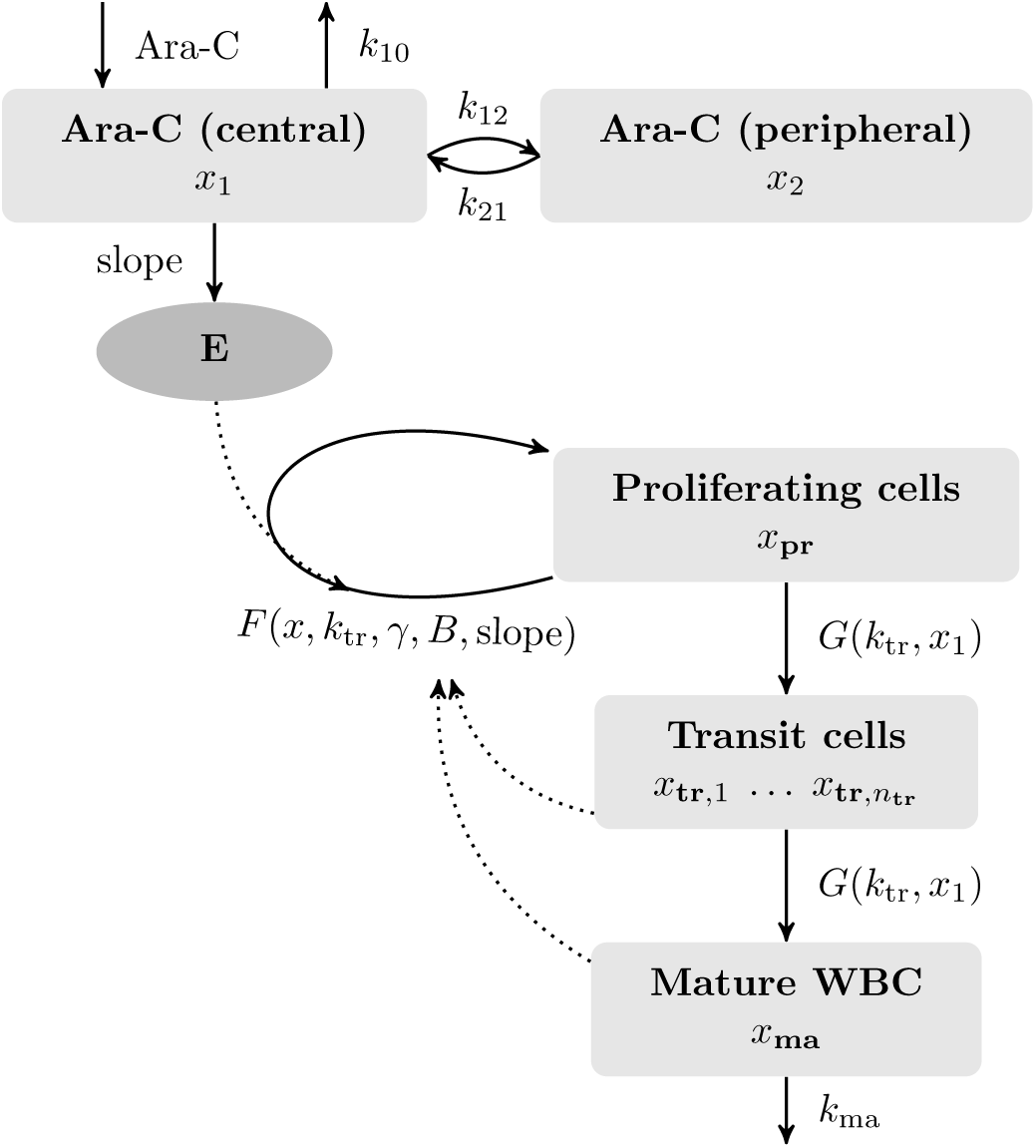
Schematic model from which all mathematical models were derived. We assumed clustering of cells and cytarabine (Ara-C) concentrations in compartments with identical properties. White blood cell (WBC) differentiation is represented by a proliferating compartment *x*_pr_, a number *n*_tr_ of transit compartments *x*_tr_ with different levels of maturation, and a compartment *x*_ma_ with mature, circulating WBC. Cells mature with a maturation rate G. Mature cells *x*_ma_ are dying by the process of apoptosis with a death rate of *k*_ma_. The pharmacodynamic effect of Ara-C is described as a log-linear function E targeting the proliferating cells in the bone marrow. It depends on the concentration *x*_1_ of Ara-C in an assumed central compartment including the circulating blood. The proliferation rate *F* of *x*_pr_ models the replication speed of proliferating progenitor cells. Modelling assumptions were incorporated by choosing different functions *F* and *G* (compare **Table 1**). The estimated model parameters used for personalisation were *B*, slope, *k*_tr_, γ, and initial conditions.

**Table 1.** Overview of all investigated mathematical models M1–M12. For each mathematical model the number of transition compartments *n*_tr_, the initial condition strategy, and the two functions *F* for proliferation rate and *G* for maturation rate are specified, compare section **Methods** and **Fig 1**, respectively. The models M1–M5 have been used mainly to determine the best number of transition compartments and initial condition strategy, which have been kept fixed from M5 onward. Different modelling assumptions are incorporated via different functions *F* and *G* in the models M5–M12. An important role has the function *S*(*x*_1_) := 1 + ln (1 + *c_V_ x*_1_), compare section **Methods**. Most important for this paper are the gold-standard model M5 which serves as a reference, and the extended models M10 and M12 as the most promising new candidate models.

| Model | $n_{\text{tr}}$ | Initial condition | Proliferation rate $F(\mathbf{x}, \mathbf{k}_{\text{tr}}, \gamma, \mathbf{B}, \text{slope})$ | $G(\mathbf{k}_{\text{tr}}, \mathbf{x}_1)$ |
| --- | --- | --- | --- | --- |
| **M1** | 6 | I1 | $(1 - E) k_{\text{tr}}(B/x_{\text{ma}})^\gamma$ | $k_{\text{tr}}$ |
| Modelling hypotheses from gold-standard model [6]. |  |  |  |  |
| **M2** | 3 | I1 | $(1 - E) k_{\text{tr}}(B/x_{\text{ma}})^\gamma$ | $k_{\text{tr}}$ |
| As M1, with $n_{\text{tr}} = 3$ transition compartments. | | | | |
| **M3** | 1 | I1 | $(1 - E) k_{\text{tr}}(B/x_{\text{ma}})^\gamma$ | $k_{\text{tr}}$ |
| As M1, with $n_{\text{tr}} = 1$ transition compartments. | | | | |
| **M4** | 1 | I2 | $(1 - E) k_{\text{tr}}(B/x_{\text{ma}})^\gamma$ | $k_{\text{tr}}$ |
| As M3, but with more freedom to choose initial values. |  |  |  |  |
| **M5** | 1 | I3 | $(1 - E) k_{\text{tr}}(B/x_{\text{ma}})^\gamma$ | $k_{\text{tr}}$ |
| As M3, but with more freedom to choose initial values. |  |  |  |  |
| **M6** | 1 | I3 | $k_{\text{tr}}(B/x_{\text{ma}})^\gamma - E$ | $k_{\text{tr}}$ |
| As M5, but assuming a direct killing effect of Ara-C on the proliferating cells. |  |  |  |  |
| **M7** | 1 | I3 | $(1 - E) k_{\text{tr}}/S(x_1)(B/x_{\text{ma}})^\gamma$ | $k_{\text{tr}}/S(x_1)$ |
| As M5, but replacing $k_{\text{tr}}$ by $k_{\text{tr}}/S(x_1)$ throughout. | | | | |
| **M8** | 1 | I3 | $(1 - E) k_{\text{tr}}/S(x_1)(B/x_{\text{ma}})^\gamma$ | $k_{\text{tr}}$ |
| As M5, but replacing $F$ by $F/S(x_1)$. | | | | |
| **M9** | 1 | I3 | $(1 - E) k_{\text{tr}}/S(x_1)(B/x_{\text{ma}})^{\gamma S(x_1)}$ | $k_{\text{tr}}$ |
| As M8, but also multiplying $\gamma$ with $S(x_1)$. | | | | |
| **M10** | 1 | I3 | $(1 - E) k_{\text{tr}}(B/x_{\text{ma}})^{\gamma S(x_1)}$ | $k_{\text{tr}}$ |
| As M5, but multiplying $\gamma$ with $S(x_1)$, possibly via macrophage activation. | | | | |
| **M11** | 1 | I3 | $(1 - E) k_{\text{tr}}(B_{\text{bm}}/(0.01 * x_{\text{pr}} + 0.99 * x_{\text{tr}}))^\gamma$ | $k_{\text{tr}}$ |
| As M5, but feedback depends on bone marrow precursor WBC instead of WBC. |  |  |  |  |
| **M12** | 1 | I3 | $(1 - E) k_{\text{tr}}(B_{\text{bm}}/(0.01 * x_{\text{pr}} + 0.99 * x_{\text{tr}}))^{\gamma S(x_1)}$ | $k_{\text{tr}}$ |
| Combining both modelling assumptions of M10 and M11. |  |  |  |  |

### M1–M5: the basic PK/PD model, number of compartments and initial conditions

In 2002 Friberg *et al*. published a PK/PD model describing myelosuppression induced by different chemotherapeutic agents (docetaxel, paclitaxel, and etoposide) [6]. The model showed a good trade-off between capturing the important aspects of the dynamics, containing a moderate number of identifiable model parameters, and being applicable for different cytostatic drugs. It has become the gold-standard model in this field with different PK and population-based modifications to topotecan [21], to daunorubicin [22], to a combination therapy of Ara-C (low-dose), etoposide and daunorubicin in the induction treatment for AML [8], to a physiologically based PK model for the induction therapy of AML patients with daunorubicin and Ara-C (low-dose) [23], to a combination therapy of carboplatin, etoposide and thiotepa [24], to paclitaxel [10], to an individual-based approach [25], and to drug specific optimisations [9]. The model assumes a clustering of cells in compartments with identical properties. WBC differentiation is represented by a proliferating compartment *x*_pr_, a number *n*_tr_ of transit compartments *x*_tr_ with different levels of maturation, and a compartment *x*_ma_ with mature, circulating WBC. Cells mature with a maturation rate constant *G* = *k*_tr_. Mature cells *x*_ma_ are dying by the process of apoptosis with a death rate constant *k*_ma_. As Monte Carlo simulations were not very sensitive, we fixed *k*_ma_ to a constant value as previously proposed [7].

Apart from different PK models which were linked to the myelosuppression model, also modifications of the structural model were proposed [10, 11]. Both models have a more detailed description of the stem cell compartment. The model from Henrich *et al*. covers a consecutive decrease of the leukocyte’s nadir in the treatment cycles achieved by a prior additional compartment mimicking the slow replication of pluripotent stem cells in the bone marrow. Mangas-Sanjuan *et al*. models a cell-cycle occuring in the bone marrow compartment covering quiescent cells which do not enter the proliferation process and are not sensitive to the pharmacodynamic effect of the treatment.

We used a two-compartment PK model of high-dose Ara-C, which is administered in the consolidation phase, with zero-order input and linear elimination based on published drug concentration-time data [26]. The corresponding parameters *k*_10_, *k*_12_, *k*_21_, *V_C_* were estimated and defined as constants for all further computations. A two-compartment PK model representing a central and peripheral compartment, see **Fig 1**, adequately described the concentration-time data and coincides with the derived values for clearance and the rate constant *k*_10_ from Table 6 [26]. Compared to a published two-compartment PK model for low-dose Ara-C in the induction treatment for AML [8] our clearance and central volume estimates are 40–50% lower, but within the inter-individual variability (IIV) [8], compare the **Appendix S1**. Due to the fact that the drug concentration-time data from Figure 2 [26] is not assigned to the patients, the IIV on the clearance and on the central volume from [8] was used for and applied on our PK model to analyse the effect of the PK variability on the different modelling hypotheses in the population approach. A more detailed discussion of the PK model and the comparison with two recently published PK models [8, 27] can be found in the **Appendix S1**. The body surface area BSA and the chemotherapy treatment *u*/duration were individually fixed by clinical procedure. The PD linking the PK model to the myelosuppression model was modelled by a log-linear function [11] *E* = slope ln (1 + *c_V_x*_1_), using the parameter slope for patient-specific calibration and chemotherapeutical effects and the constant *c_V_* for unit consistency (see **Table S3**). We also implemented a linear PD function with discouraging results. Additionally, we tested a (sigmoid) *E_max_* model without achieving better model accuracies. The function *F*(*x, k*_tr_*, γ, B*, slope) is a general description of the proliferation rate of *x*_pr_ and incorporates the PD effect *E* on the proliferating cells, as discussed in Minami *et al*. [28], Derendorf *et al*. [29] and applied, e.g. in Quartino *et al*. [7]. The basic structure of the function F is (1 − *E*) *k*_tr_(*B/x*_ma_)*^γ^* in which the mature cells influence the proliferation rate *k*_tr_ of *x*_pr_ with a feedback term (*B/x*_ma_)*^γ^* that leads to higher rates if the number of circulating cells *x*_ma_ is below the baseline WBC count *B*, and vice versa. The proliferation exponent *γ* indicates the strength or speed of this feedback. The estimation parameters were *B*, slope, *k*_tr_, and *γ* plus a varying approach of initial conditions. We implemented three different strategies to treat initial conditions: I1 assumes a steady state, I2 assumes a steady state only for the proliferating and transient compartments, and I3 penalises deviations from the steady state. I2 and I3 are considered as alternative initial conditions as our clinical data indicate that the steady state assumption after induction phase and between the consolidation cycles may be violated. These two approaches may also account for disease progression effects. A specification of I1, I2, I3 and a discussion of the number of compartments can be found in the **Appendix S1**. We used I1 and *n*_tr_ = 6 for M1, I1 and *n*_tr_ = 3 for M2, I1 and *n*_tr_ = 1 for M3, I2 and *n*_tr_ = 1 for M4, and I3 and *n*_tr_ = 1 for M5–M12.

**Fig 2.**
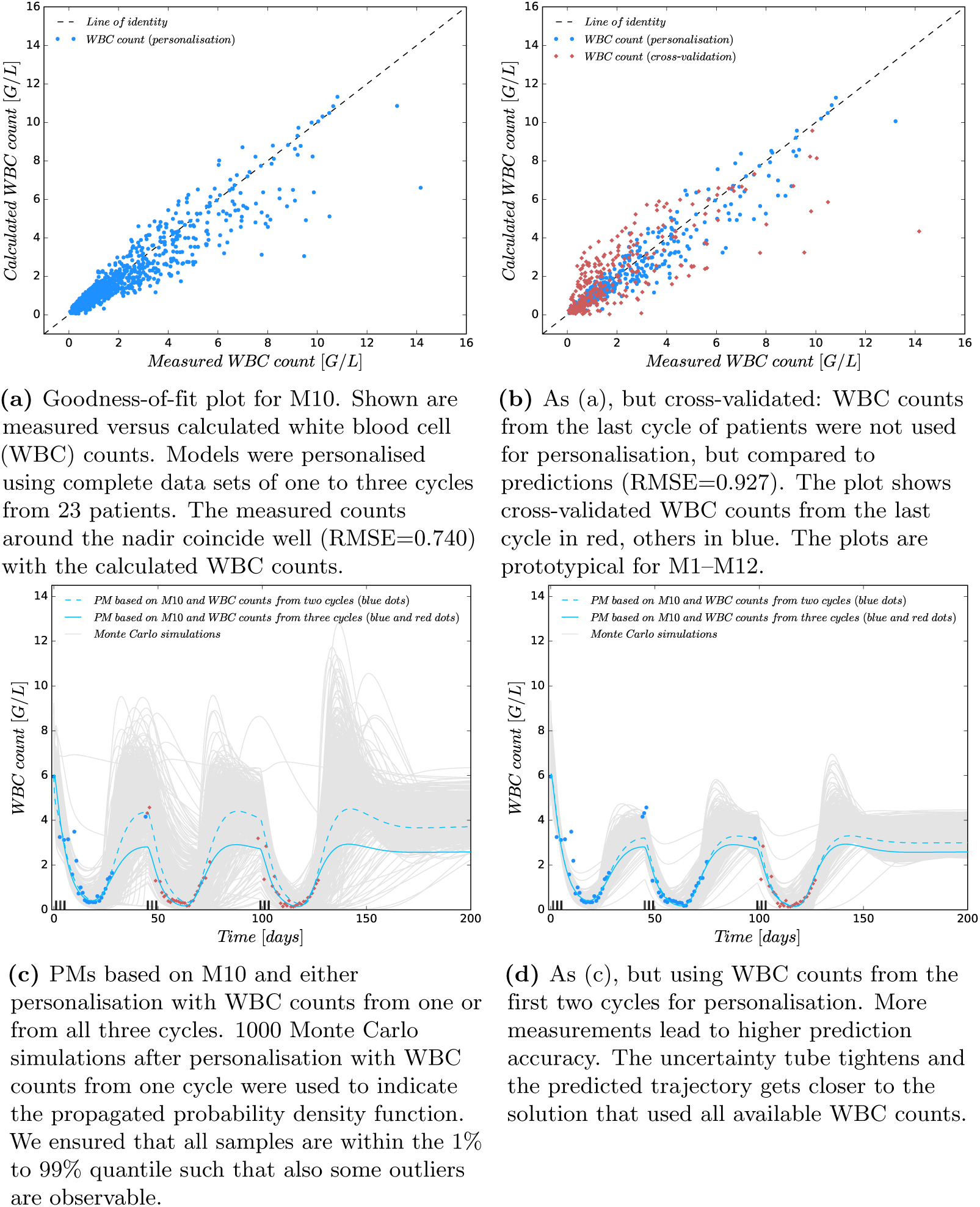
Visualisation of predictive accuracies of personalised mathematical models (PM).

### M6: Modelling a Direct Killing Effect of Ara-C on the Proliferating Cells

In the model M6, we chose the proliferation rate as discussed in previous works [9, 23, 29] as *F* = *k*_tr_(*B/x*_ma_)^γ^ − *E*. The main difference to all other models is that the PD effect *E* is directly multiplied with *x*_pr_ and not with *k*_tr_(*B*/*x*_ma_)^γ^*x*_pr_. Multiplying with *x*_pr_ can be seen as a direct (killing) impact of Ara-C on the amount of proliferating cells, whereas the more plausible mechanism-based rationale is the induced reduction of the proliferation rate constant *k*_tr_ used in all models except in M6.

### M7–M12: Extending the Effects of Ara-C

The root mean squared error (RMSE) values in **Table 2** indicate that model M5 with one transition compartment and initial condition approach I3 provides the highest accuracy after model personalisation compared to M1–M4.

**Table 2.** Root mean squared error (RMSE) values for the models M1–M12. Measured and calculated WBC counts were compared. The estimations and predictions used personalised mathematical models (PMs) that were calculated based on the twelve different mathematical models M1–M12. The first row refers to a personalisation for all 42 consolidation cycles (CCs). The second row shows results for personalisations using all available cycles per patient (Pat). For predictions (Pred) all but one cycle were used for personalisation and the last cycle for cross-validation. Four more rows show the predictions seperated into the different schedules (D135, d135, *P*_D123_ and *P*_D12_). The RMSE values decrease from cycles to patients and from personalisation towards prediction, as expected. Comparing the mathematical models, the accuracy increases with a reduced number of compartments from M1 to M3. The initial condition strategies I2 in M4 and I3 in M5 decrease RMSEs further. M5–M12 all used *n*_tr_ = 1 and I3 and performed equally well, with the slight exception of M7. Note that in particular there is no significant difference between the established gold-standard model M5 and our newly proposed extended model M10.

| RMSE | M1 | M2 | M3 | M4 | M5 | M6 | M7 | M8 | M9 | M10 | M11 | M12 |
| --- | --- | --- | --- | --- | --- | --- | --- | --- | --- | --- | --- | --- |
| 42 CCs | 0.911 | 0.836 | 0.742 | 0.636 | 0.579 | 0.595 | 0.639 | 0.576 | 0.574 | 0.574 | 0.577 | 0.587 |
| 23 Pat | 1.154 | 1.011 | 0.892 | 0.825 | 0.741 | 0.758 | 0.785 | 0.753 | 0.738 | 0.740 | 0.740 | 0.741 |
| 14 Pred | 1.269 | 1.128 | 1.059 | 1.007 | 0.908 | 0.972 | 0.997 | 0.960 | 0.958 | 0.927 | 0.940 | 0.947 |
| 7 D135 | 1.108 | 0.912 | 0.834 | 0.778 | 0.750 | 0.753 | 0.781 | 0.750 | 0.767 | 0.765 | 0.731 | 0.768 |
| 5 d135 | 1.319 | 1.240 | 1.141 | 1.093 | 0.921 | 1.095 | 1.023 | 1.068 | 1.043 | 0.957 | 1.037 | 1.009 |
| $P_{D123}$ | 2.404 | 2.218 | 2.241 | 2.258 | 2.011 | 2.014 | 2.324 | 2.006 | 1.996 | 1.996 | 2.029 | 2.022 |
| $P_{D12}$ | 1.014 | 0.991 | 1.042 | 0.924 | 0.842 | 0.843 | 1.049 | 0.839 | 0.840 | 0.843 | 0.824 | 0.823 |

The indirect effect of Ara-C with an impaired proliferation (M5) is more plausible than a direct killing effect (M6), because Ara-CTP is incorporated into DNA and RNA and impairs cell replication [14]. Therefore, M5 became the reference model for all further analysis. We extended the proliferation rate *F*(·) and/or the transition rate *G*(·) in M5 to capture potential secondary effects of Ara-C. To understand the implications of the extensions, we observe that the proliferation rate *F* = (1 − *E*) *k*_tr_(*B*/*x*_ma_)*^γ^* is negative when 1 < *E*. This is the case for

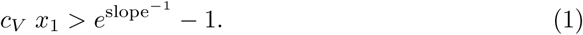

This corresponds to more proliferating cells being in the process of apoptosis than being in the process of cell division. It is important that the feedback term (*B*/*x*_ma_)*^γ^* increases the absolute value of *F* for *B* > *x*_ma_, and decreases it for *B* < *x*_ma_. Therefore, an analysis of *F* always has to consider all four cases related to the signs of 1 − *E* and of *B* − *x*_ma_. Inspired by the log-linear behaviour of the PD effect *E*, we chose

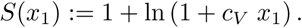

This monotonously increasing function is applied to different expressions in M5.

In M7 we replaced the transition rate *k*_tr_ by *k*_tr_/*S*(*x*_1_) throughout M5. This results in an Ara-C induced reduction of the transition rate.

In M8 we replaced the complete feedback function *F* in M5 by *F*/*S*(*x*_1_). This models an Ara-C induced decreased auto-feedback of the proliferating cells. For high values of *x*_1_, i.e. when (1) holds, this results in a decreased killing of proliferative cells. For values *x*_1_ > 0 below that boundary, we get a decreased positive proliferation rate.

In M9 we replaced both the complete feedback function *F* by *F/S*(*x*_1_) and the proliferation exponent γ by γ *S*(*x*_1_). Again, depending on *x*_1_ either the killing or the proliferation rate of *x*_pr_ are decreased by *F/S*(*x*_1_). In addition, the impact depends on whether the WBC count is below or above the baseline: for *x*_ma_ *< B* we have an increased killing/proliferation rate 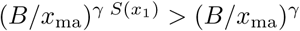 and vice versa.

In M10 we replaced the proliferation exponent γ in M5 by γ *S*(*x*_1_). This is motivated by the observation that the feedback term with exponent γ is related to the endogenous G-CSF [6], and the assumption that higher concentrations of Ara-C coincide with activation of macrophages and hence increased G-CSF secretion. In contrast to M9 the function F itself is not scaled. Like in M9, the γ *S*(*x*_1_) scaling results in an increase of killing/proliferation rates for WBC counts below the baseline, and a decrease else.

In M11 we replaced the quotient *B*/*x*_ma_ by a comparison between cells in the bone marrow and their baseline value. We estimated that about 1% of the WBC precursor cells in the bone marrow are in the proliferating compartment *x*_pr_, and 99% in the transition compartment *x*_tr_.

In M12 we combined the extensions from M10 and M11.

### Clinical Data (high density WBC counts) & Personalisation

AML patients who had received induction therapy (commonly defined as anthracycline- and Ara-C-based *7+3 regimen* [2]) resulting in complete remission and who did not receive granulocyte-colony stimulating factors (G-CSF) during the post-remission consolidation therapy were eligible for data analysis. We focused on patients who did not receive growth factor support, as such effects were not yet accounted for in our mathematical models. Almost daily WBC counts from 42 consolidation Ara-C cycles (CCs) of 23 AML patients (median 62 years, 14 male, mostly *de novo* AML (19/23), mostly AML FAB-M2 (9/19), mostly intermediate cytogenetic risk (12/20)) from 2008 to 2015 were analysed from clinical charts provided by the Department of Hematology and Oncology, Magdeburg University Hospital, Magdeburg, Germany. The data were retrospectively collected and pseudonymised from records of the clinical routine. Interventions were not performed for this work. All clinical procedures were performed in accordance with the general ethical principles outlined in the Declaration of Helsinki. For this reason no patients’ agreements were required. The CCs were partitioned in one, two, and three consecutive CCs from nine, nine, and five patients, respectively. Four different schedules D135, d135, D123, or D12, in which the numbers correspond to treatment days 1, 2, 3, and 5, respectively, d to intermediate-dose Ara-C (i.e. 1 *g/m*^2^ per body surface area (BSA) twice a day over three hours) and D to high-dose Ara-C (i.e. 3 *g/m*^2^ twice a day), were administered 24, 14, two, and two times. Patient *P*_D123_ (62 years, male) received two cycles of D123. Patient *P*_D12_ (64 years, female) received two cycles of D12. The 21 other patients received 1-3 D135 cycles (median 57 years, 8 male, 4 female) or d135 cycles (median 68 years, 5 male, 4 female).

We used all 42 CCs to personalise our mathematical models M1–M12 performing point estimations (individual approach) and M3, M10 (with I1), the model from Henrich *et al*. [10] and the model from Mangas-Sanjuan *et al*. [11] applying nonlinear mixed-effects modelling (population approach).

The ordinary differential equations comprise states (drug concentrations and cell counts), control (chemotherapy schedule), and model parameters. Model parameters are variable numbers and the main tool for personalisation. In the interest of identifiability we calculated values for PK parameters a priori from published data [26] and fixed others to values from the literature. We then used individual and population parameter estimations to obtain the remaining model parameters *B*, slope, *k*_tr_, and γ plus a varying initial condition approach such that measured WBC counts from one or several cycles were fitted in an optimal way. Once the model parameters have particular values, the model is called PM. A PM can be solved numerically, resulting in calculated cell counts at arbitrary time points that can be used for further analysis.

### Prediction & Cross-Validation

The PMs were then used to predict (simulate) and cross-validate WBC counts for the last CC of 14 patients for whom at least two consecutive CCs are available. Additionally, we calculated predicted *t*_rec_ values from our 42 PMs (see **Clinical Data & Personalisation**) applying D123 and D135 schedules and compared the descriptive statistics with published average *t*_rec_ values from a subset of data (367 CCs of 208 AML patients, no G-CSF support) of the AMLSG 07-04 trial in which the schedules D123 and D135 after *7+3 regimen* were analysed [30]. The published AMLSG 07-04 [30] trial does not provide WBC counts to obtain new PMs, therefore we used the median of observed *t*_rec_ values for D123 and D135 Ara-C schedules. In the interest of a fair comparison (i.e., to avoid comparison with the value 0) we excluded five out of 42 PMs for which at least one prediction (M1–M12 with either D123 or D135) resulted in no WBC counts below the threshold value. Further, we predicted *t*_rec_ values for two Ara-C schedules in which a constant administration of Ara-C throughout days 1-5, with either 100 *mg/m*^2^ or 400 *mg/m*^2^ was given. These schedules, together with D135, have been clinically analysed for 1088 AML patients (median 52, 568 male) by Mayer *et al*. [3], and the superiority of D135 with respect to disease-free survival rates and remaining in continuous complete remission after four years has been shown but no *t*_rec_ values were reported. Finally, we analysed the effect of the inter-individual PK variability on the *t*_rec_ values derived by the models M3 and M10 (with I1). We applied schedules D123 and D135 with fixed population parameter values for *B*, slope, *k*_tr_, and γ and performed 500 simulations each with randomly chosen values from the inter-individual variability (IIV) for the PK parameters clearance and central volume.

All experiments were performed to analyse the 12 proposed models with respect to WBC count and *t*_rec_ predictability.

### Schedule Timing

After verifying the predictability performance of the PMs, we performed a simulation study in which we demonstrated a further possible application of the PMs in clinical practice. We analysed the impact of the treatment timing on the individual nadir values. For each of the 14 patients, for whom at least two consecutive CCs were available, the nadir of the last CC was compared to 20 simulated nadirs. These nadirs resulted from simulations using the patient’s PMs (second row of Table 2) in which the timing of the last CC was varied daily with the maximal starting variation of 10 days earlier or later.

## Results

### Accuracy of PMs with fixed Ara-C schedule

**Table 2** shows statistics for PMs derived from M1–M12, for a pure estimation (using all available WBC counts to personalise the model) and for a cross-validation (using all but the last CC for personalisation). The accuracies depend strongly on the number of compartments and initial condition strategy (M1–M5), but are stable against modelling uncertainty with respect to possible effects of Ara-C (RMSE values for prediction between 0.997 and 0.908 for M5–M12). These values were even better when the standard schedule D135 was applied in the estimated and predicted cycles. Goodness-of-fit plots in **Fig 2a**−b and **Fig S1** visualise the good match between model predictions and measured WBC counts (respectively observed *t_rec_* values) around the nadir and a wider spread of large WBC counts. **Fig 2c**−d indicate the involved uncertainty derived from the individual variance-covariance matrix by means of Monte Carlo simulations (for more information see **Appendix S1**). The uncertainty reduces when more WBC counts are used, and the accuracy increases.

The WBC counts around the nadir are explained well by all models for fixed Ara-C schedules (either D135 or D123), as shown in **Fig 3a**–**d** for two exemplary patients and in **Fig S2-S3** for the other 12 patients with at least two consecutive CCs.

**Fig 3.**
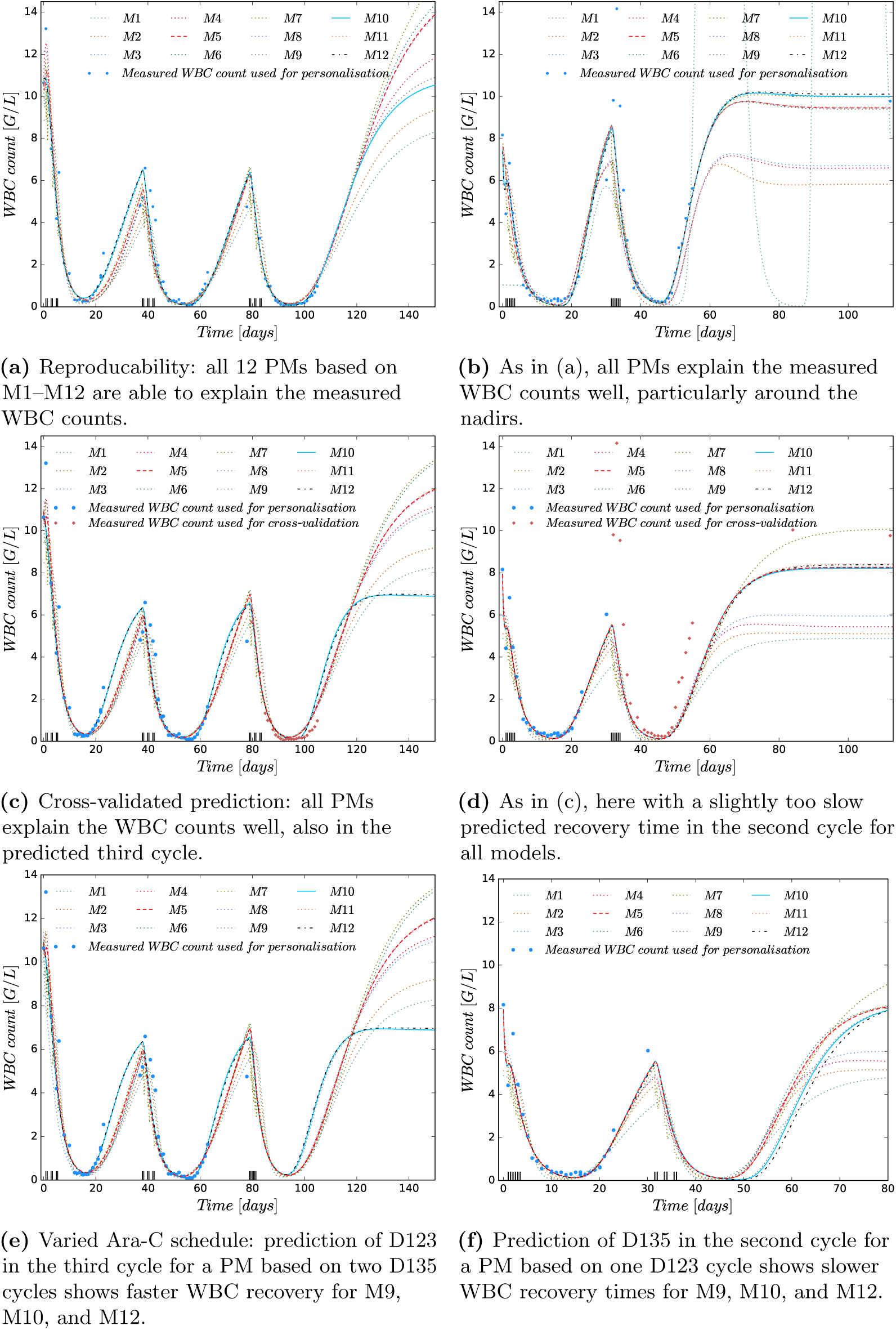
Comparison of personalised models (PMs) based on M1-M12 and white blood cell (WBC) data. Patient with three D135 cycles (left) and patient *P*_D123_ with two D123 cycles (right), as indicated on the x-axis. The PMs exemplify reproducability (first row), predictability (second row) and simulation of a different schedule in prediction than estimation (third row).

### Accuracy of PMs with altered Ara-C schedule

**Fig 3e**–f show two cases where D135 is used for personalisation and D123 for prediction (and vice versa). Here, M9, M10, and M12 have a faster (slower) hematological recovery for D123 (D135). All three models assume that the proliferation speed γ depends on the Ara-C concentration. This modelling assumption is visualised in a different way for M5, M10 and M12 in **Fig S4–S6**. It is shown that the proliferation rate *F* of M10 and M12 compared to the proliferation rate of M5 behaves in such a way that a faster WBC recovery for D123 schedules is achieved. This indicates that the comparison of WBC recovery times between D123 and D135 treatments is a suitable criterium for model discrimination.

**Table 3** shows results for a comparison of predicted and measured *t*_rec_ values. We used 444 PMs (using M1–M12 and clinical data from 37 cycles with schedules D135, d135, D123 and D12 from section **Clinical Data & Personalisation**) to predict the outcome of D135 and D123 schedules. The median values of the predicted *t*_rec_ were compared to the values from a subset of data (108 with D135 and 259 with D123 schedules) from the AMLSG 07-04 trial [30]. M9, M10, and M12 resulted in roughly 4 days faster *t*_rec_ for D123 compared to D135, similar to the clinical result from the literature and in contrast to the 1 day difference of M5. The individual results have been qualitatively confirmed by the predicted *t*_rec_ values from the population approach (see **Table S4**). The models from Henrich *et al*. and Mangas-Sanjuan *et al*. were not further considered, as both models simplified to the original Friberg model after parameter estimation. For the model from Henrich *et al*., the estimated population parameter value *f_tr_* was 0.96, supporting the visual assessment that the patients’ nadirs are not decreasing during the CCs. The estimated parameter values *k_cycle_* = 0.0009 and *F_prol_* = 0.941 of the model from Mangas-Sanjuan *et al*. yielded a non-existing stem cell cycle. A possible reason for the non-identifiability of the parameters might be the limited schedule variation. The authors state that a vast variation of schedules has to be available for parameter identification [11].

**Table 3.**
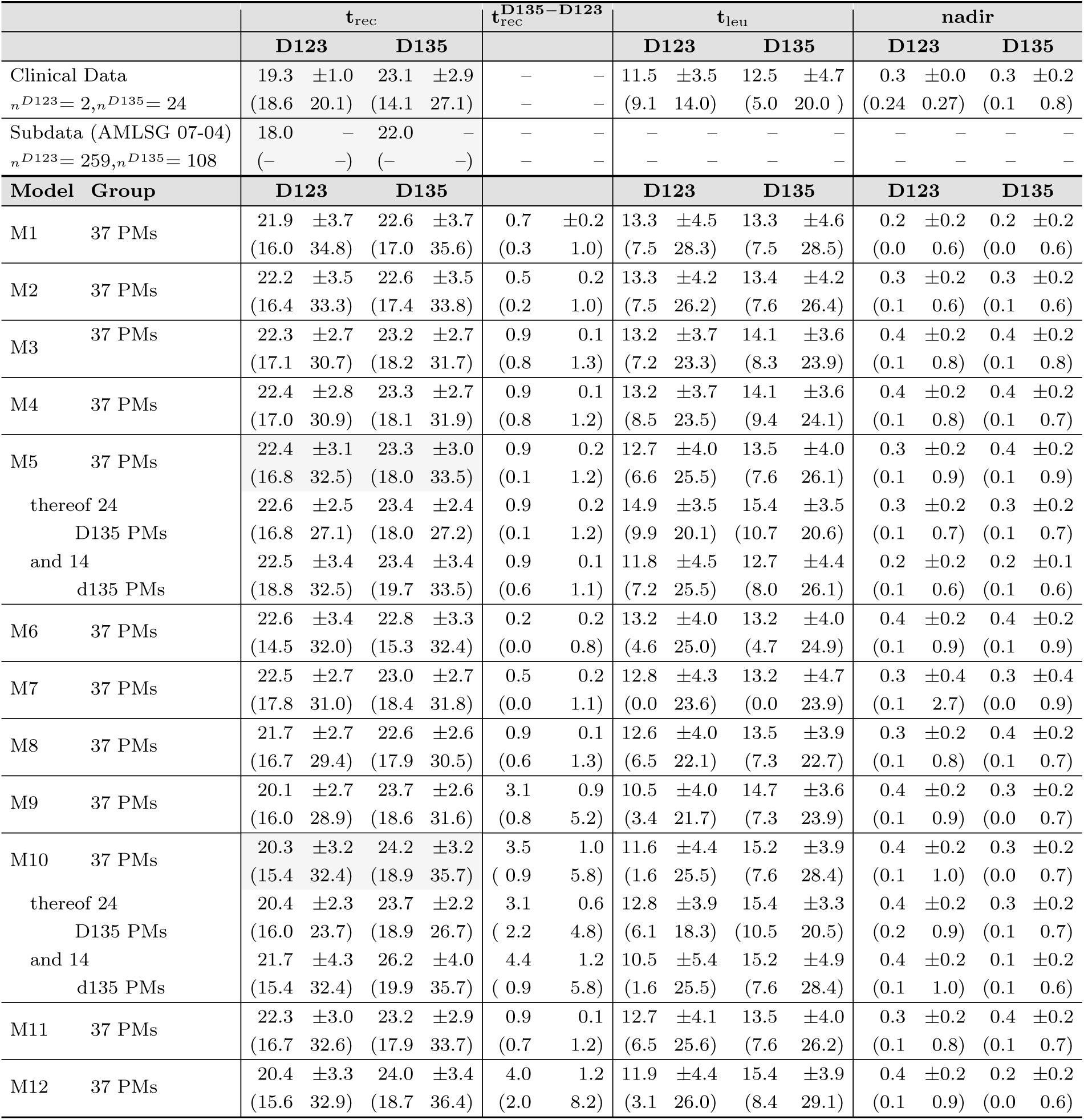
Double cross-validation with clinical data from two independent clinical trials. Shown are the median, standard deviation, minimum and maximum (in brackets) of *t*_rec_, the *leukopenia time t*_leu_ (the number of days with WBC count ≤ 1 *G/L)* and *nadir* for D123 and D135 schedules. The first two rows show values from two independent clinical studies that serve as a comparison. The second part of the table shows prediction results. Predictions were calculated with PMs from our clinical data with underlying mathematical models M1–M12. Model M5 explained well the outcome of schedule D135, but showed a significant mismatch of more than three days for schedule D123. The predictions using the extended model M10 were better for schedule D123. See also **Fig 3e**–**f** for an illustrated comparison between M5 and M10. Model M9 and M12 were also promising, but we focused on M10 applying Ockam’s razor.

The simulation study analysing the effect of the PK variability on the resulting recovery times of schedules D123 and D135 for models M3 and M10 (with I1) revealed that model M10 was more sensitive to different high-dose Ara-C treatment schedules compared to model M3 despite the high inter-individual PK variability. This was verified in **Fig S9** presenting boxplots of 500 simulated *t*_rec_ values for both models and schedules with IIV on the PK.

### Schedule Timing

We analysed the impact of different treatment starts of the last CCs with respect to subsequent nadir values. A comparison to the clinically observed nadir values indicated a large potential for clinical improvement, i.e., a higher nadir value due to a different treatment timing (see **Fig 4(a)**). **Fig 4(b)** exemplarily shows the WBC dynamics for different treatment timings. Earlier (later) starts resulted in sequentially higher (lower) nadir values.

**Fig 4.**
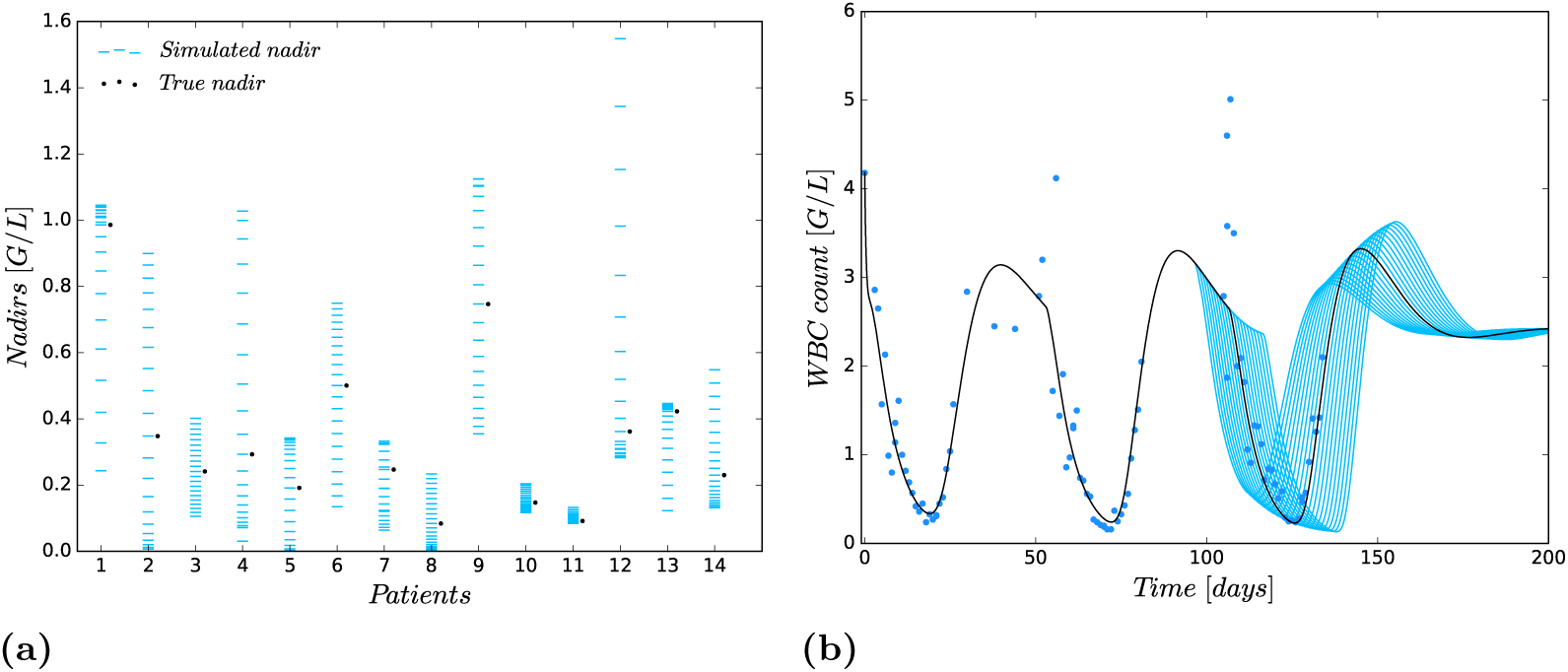
Analysing the influence of treatment timing on nadir values. **(a)** Simulation study in which 20 simulated nadirs were compared with the true nadir of the last CC for the 14 patients who have more than one CC. The simulated nadirs were computed by using the patient’s PM (second row of Table 2) and varying the start of the last CC daily with the maximal starting variation of 10 days earlier or later. **(b)** Exemplary variation of the CC start for one patient. An earlier (later) start results in a larger (lower) nadir.

## Discussion

High-density WBC counts from 23 AML patients were collected and used to personalise 12 mathematical models and analyse their prediction accuracy with respect to different modelling hypotheses and treatment schedules. The high prediction accuracies of the PMs, especially around the nadir, confirm previous claims [4, 12] that the general approach of in-silico studies can be used for clinical decision support, e.g. to monitor and predict WBC dynamics.

In combination with clinical expertise on the impact of schedules on relapse probabilities, this might have an important clinical impact via altered treatment schedules which might eventually result in improved survival rates. To evaluate or even optimise the dosage of Ara-C, PMs may not be appropriate, though. We showed that an analysis based on a fixed chemotherapy schedule can not discriminate between different modelling hypotheses. The agglomerative nature of the mathematical models leads to a choice of model parameters that is not only personalised to the patient, but also to the applied schedule. Therefore, we used, different schedules for personalisation and prediction to overcome this problem and to allow discrimination of the models. This approach facilitated to distinguish between the modelling hypotheses implemented in models M5–M12 and enabled to find the suitable model assumption considered in M9, M10, and M12. In our opinion this procedure should be routinely applied, preferably using high density WBC counts for different schedules in the same patients. As an alternative to such a tedious clinical study we suggest to use average *t*_rec_ values as a discrimination criterion for competing models.

Comparing our clinical data and the AMLSG 07-04 trial with respect to *t_rec_* in **Table 3** and **Table S4**, our observed WBC recovery times are for both schedules 1 day longer. This can be explained by the age difference between patients in our clinical data (median 62 and 57 years for D123 and D135, respectively) and the subdata of the AMLSG 07-04 trial (median of all patients in the trial 49 years) and a related statistical analysis: Jaramillo *et al*. [30] found in a multivariable analysis a significantly longer WBC recovery for older patients (hazard ratio of a 10-year age difference, 0.89; P = 0.001) [30] and a significantly shorter WBC recovery for patients receiving D123 compared to the reference group D135 (hazard ratio, 1.94; P < 0.0001) [30] which coincides well with our findings. The comparison of *t*_rec_ values for D123 and D135 treatments indicates that models M9, M10, M12 are the best candidates among M1-M12 for future work on the simulation and optimisation of intermediate to high-dose Ara-C treatment schedules.

Summarising, we extended the gold-standard model for myelosuppression [6] to the most important component in consolidation therapy [2, 13], Ara-C, and showed that one modelling assumption was important for a faster WBC count recovery for D123 schedules. In models M9, M10, and M12 we assumed that the Ara-C concentration has a direct impact on the proliferation speed. As stated above, such a modelling assumption has an agglomerative nature and the underlying physiological processes are still unknown. We speculate that a high Ara-C concentration might lead to an increased number of cell deaths and thereby induces phagocytosis and macrophage activation, which in turn might increase G-CSF secretion and hence proliferation speed. Future G-CSF concentration measurements for AML patients during consolidation cycles of D123 and D135 treatments and a comparison of our extended models with Quartino’s [31] integrated G-CSF-myelosuppression model may shed light on this speculation.

## Acknowledgements

The authors wish to thank Franziska Kluwe, Graduate Research Training Program PharMetrX and Department of Clinical Pharmacy and Biochemistry, Institute of Pharmacy, Freie Universitaet Berlin, for her support in the pharmacokinetics and nonlinear mixed-effects modelling analysis.

## Author Contributions

FJ extended and implemented the mathematical models and did all numerical computations. ES and TF contributed to modelling, study design, and provided clinical data. KR contributed to data aquisition, mathematical modelling and developed the PK model for Ara-C. SS contributed to mathematical modelling, numerical approaches, and study design. All authors contributed to discussion of results and writing of the final paper.

## Data Availability

Raw data contain patient-identifying information and are unsuitable for public deposition. Anonymized data sets can be obtained by contacting Prof. Dr. Sebastian Sager.

## Supporting Information

### Appendix S1 Additional methods

**Table S1 Terminology and potentially confusing synonyms.** Expressions that are also used in the manuscript are in *italic*.

**Table S2 Comparison of model predictions for low-dose treatment schedules.** As in **Table 3** predicted values for different treatment schedules are shown, based on underlying mathematical models M1–M12. Shown are the values of median, standard deviation, minimum and maximum (in brackets) for two low-dose schedules. Both assume a constant administration of drugs throughout days 1 to 5, with either 100 *mg/m*^2^ or 400 *mg/m*^2^ body surface. We do not have clinical data to compare these predictions, but they give additional insight on the possibility to discriminate models M1–M12 and a general trend showing that independent of the modelling assumptions both low-dose schedules result in slower white blood cell recovery and lower nadirs than D135 or D123. This has been clinically observed [3]. (Two further personalised cycles were excluded because for some models no recovery after chemotherapy was observed)

**Table S3 Model constants, patient-specific constants, and units of model parameters.** The values were used to obtain personalised mathematical models. The constants were determined from published data [26] and applied to all patients. To shorten notation we also used 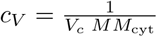. The patient-specific infusion times and dosages that define a treatment schedule were modified for simulation and optimisation of different schedules. The range shows minimum and maximum values of all considered data in the clinical study.

**Table S4 Objectives (final objective function values from FOCEi method (OBJ), population predicted 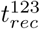 and 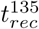 values), parameter and coefficient of variation (CV) estimates with relative standard errors (RSE) from nonlinear mixed-effects modelling of models M3 and M10 with initial condition approach I1 (with I1).**

**Figure S1 Goodness-of-fit plot for all but three (because of WBC counts greater 1) measured and calculated *t_rec_* values for model M10.** The measured *t*_rec_ values are slightly higher due to the coarser measurement grid.

**Figure S2 Cross-validation of predicted white blood cell (WBC) counts from personalised models (PMs) M1-M12 and measured WBC counts for six patients treated with D135.** All but one cycle were used for personalisation and the last cycle for cross-validation. For patients (a)-(e) the PMs can predict the WBC count decrease after Ara-C administration in the last cycle where models M10 and M12 have a slower WBC recovery than M5. For patient (f) the WBC recovery from the PMs starts to early compared to the measured WBC counts.

**Figure S3 Cross-validation of predicted white blood cell (WBC) counts from personalised models (PMs) M1-M12 and measured WBC counts for five patients (a)-(e) treated with d135 and one patient (f) treated with D12.** The PMs provide good predictions for patient (a) and (f) but show mismatches in recovery times and nadir values for patients (b)-(e).

**Figure S4 Comparing personalised mathematical models (PMs) M5 and M10 for D123 and D135 schedules (data set I).**

**Figure S5 Comparing personalised mathematical models (PMs) M5 and M10 for D123 and D135 schedules (data set II).**

**Figure S6 Comparing personalised mathematical models (PMs) M10 and M12 for D123 and D135 schedules (data set I).**

**Figure S7 Simulations of different pharmacokinetic models, Ara-C concentration measurements and inter-individual variability.**

**Figure S8 Visual predictive checks (VPCs), derived by 1000 simulations, for leukocytes** [*G/L*] **versus time** [*days*] **starting with the first measurement before dosing for model M3 (a) and M10 (with I1) (b)**. Blue circles are the measured WBC counts of 23 AML patients described in section **Clinical Data & Personalisation**. One measurement was taken at timepoint 88.98 [days] with the value 7.18[G/L] which is not shown in the VPCs. Red lines show the median (solid) and 5th and 95th percentiles (dashed) of measurements. The shaded areas represent the 90% confidence intervals around the 5th (blue), 50th (red) and 95th (blue) simulated percentiles of the model predictions. Regarding the VPCs, model M3 and M10 have an almost equivalent prediction accuracy. The 50% percentiles of measurements and model predictions perfectly overlap, thus supporting our individually based results from **Table 2**. The same applies to the start of the 5% and 95% percentiles until the nadir. After the nadir the 5% and 95% percentiles of the model predictions recover slightly faster/slower compared to the measurements. At day 30 the percentiles of measurements and model predictions coincide again.

**Figure S9 Simulations of different pharmacokinetic models, Ara-C concentration measurements and inter-individual variability.**

